# Consensus on Molecular Subtypes of Ovarian Cancer

**DOI:** 10.1101/162685

**Authors:** Gregory M Chen, Lavanya Kannan, Ludwig Geistlinger, Victor Kofia, Zhaleh Safikhani, Deena M A Gendoo, Giovanni Parmigiani, Michael Birrer, Benjamin Haibe-Kains, Levi Waldron

## Abstract

**INTRODUCTION:** Various computational methods for gene expression-based subtyping of high-grade serous (HGS) ovarian cancer have been proposed. This resulted in the identification of molecular subtypes that are based on different datasets and were differentially validated, making it difficult to achieve consensus on which definitions to use in follow-up studies. We assess three major subtype classifiers for their robustness and association to outcome by a meta-analysis of publicly available expression data, and provide a classifier that represents their consensus.

**METHODS:** We use a compendium of 15 microarray datasets consisting of 1,774 HGS ovarian tumors to assess 1) concordance between published subtyping algorithms, 2) robustness of those algorithms to re-clustering across datasets, and 3) association of subtypes with overall survival. A consensus classifier is trained on concordantly classified samples, and validated by leave-one-dataset-out validation.

**RESULTS:** Each subtyping classifier identified subsets significantly differing in overall survival, but were not robust to re-fitting in independent datasets and grouped only approximately one third of patients concordantly into four subtypes. We propose a consensus classifier to identify the minority of unambiguously classifiable tumors across multiple gene expression platforms, using a 100-gene signature. The resulting consensus subtypes correlate with patient age, survival, tumor purity, and lymphocyte infiltration.

**CONCLUSIONS:** Our analysis demonstrates that most HGS ovarian cancers are not able to be subtyped. A minority of tumors can be classified and our proposed consensus classifier consolidates and improves on the robustness of three previously proposed subtype classifiers. It provides reliable stratification of patients with HGS ovarian tumors of clearly defined subtype, and will assist in studying the role of polyclonality in the majority of tumors that are not robustly classifiable.

## Introduction

Ovarian cancer is a genomically complex disease, for which the accurate characterization of molecular subtypes is difficult but is anticipated to improve treatment and clinical outcome^1^. Substantial effort has been devoted to characterize molecularly distinct subtypes of high-grade serous (HGS) ovarian cancer (Table 1). Initial large scale efforts to classify HGSC of the ovary did not reveal any reproducible subtypes^2^. Tothill *et al^3^* reported four distinct HGS subtypes: (*i*) an immunoreactive expression subtype associated with infiltration of immune cells, (*ii*) a low stromal expression subtype with high levels of circulating CA125, (*iii*) a poor prognosis subtype displaying strong stromal response, correlating with extensive desmoplasia, and (*iv*) a mesenchymal subtype with high expression of N/P-cadherins. The Cancer Genome Atlas (TCGA) project also identified four subtypes characterized by (*i*) chemokine expression in the immunoreactive subtype, (*ii*) proliferation marker expression in the proliferative subtype, (*iii*) ovarian tumor marker expression in the differentiated subtype, and (*iv*) expression of markers suggestive of increased stromal components in the mesenchymal subtype, but did not report differences in patient survival^4^. Further experimental characterization revealed an increased number of samples with infiltrating T lymphocytes for the immunoreactive subtype, whereas desmoplasia, associated with infiltrating stromal cells, was found more often for the mesenchymal subtype^5^. Konecny *et al*.^6^, independently evaluated the TCGA subtypes and also reported the presence of the four transcriptional subtypes using a *de novo* clustering and classification method.

**Table 1:**
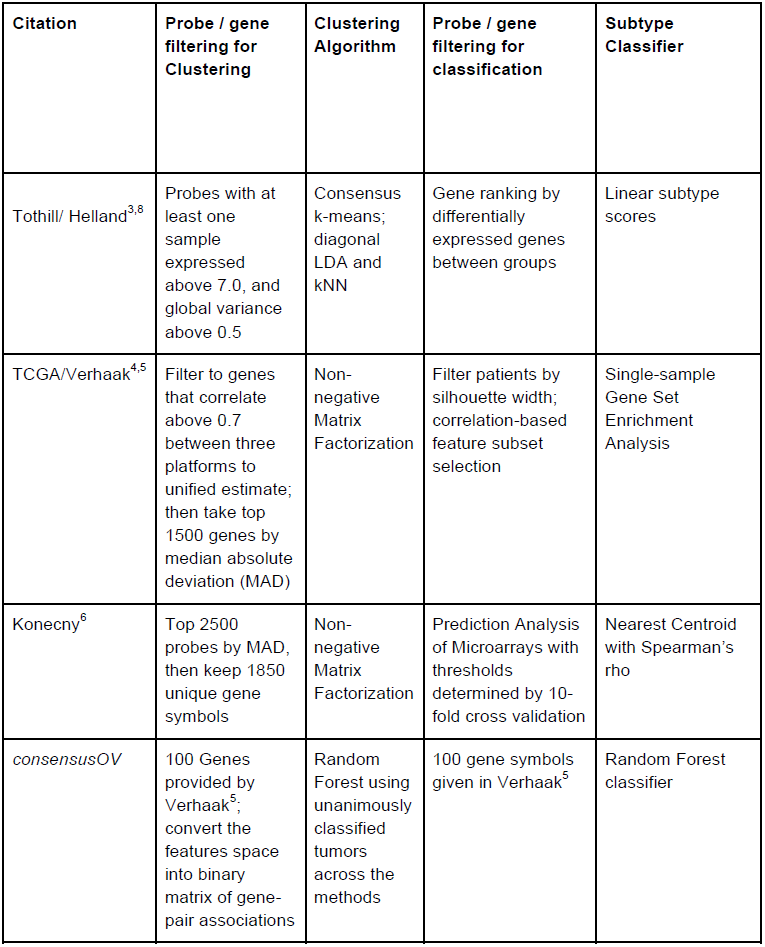
***Subtyping methodology of the algorithms reviewed.***

However, robustness and clinical relevance of these subtypes remain controversial^7^. The previous subtyping efforts have assessed prognostic significance in different patient cohorts, and have taken different approaches to validate these subtypes in independent datasets. A recent review of ovarian cancer subtyping schemes highlighted the difficulty of comparing results of studies that used different subtyping algorithms, and that better general agreement on how molecular subtypes are defined would allow more widespread use of expression data in clinical trial design.^1^ ^364^

Assessing the generalizability of subtyping algorithms is challenging as true subtype classifications remain unknown. This challenge is evident in the lack of published validation of the proposed HGS subtypes. Subsequent efforts have performed *de novo* clustering of new datasets and noted similarity in the clusters identified, but they have not reported quantitative measures such as classification accuracy or rate of concordance with previously published algorithms. In this article, we address these limitations by re-implementing three major subtyping methods^3,5,8^ and assess between-classifier concordance and across-dataset robustness. We show that these subtype classifiers yield significant concordance, and are virtually identical for tumors classified with high certainty. Using the core set of tumors concordantly classified by each method, we develop *consensusOV*, a consensus classifier that has high concordance with the three classifiers, therefore providing a standardized classification scheme for clinical applications.

## Materials and Methods

### Datasets

Analysis was carried out on datasets from the curatedOvarianData compendium9. Datasets were additionally processed using the MetaGxOvarian package10 (Supplementary Information). Analysis was restricted to datasets featuring microarray-based whole-transcriptome studies of at least 40 patients with late stage, high-grade, primary tumors of serous histology. This resulted in 15 microarray studies, providing data for 1,774 patients (Supplementary Table 1). Duplicated samples identified by the doppelgangR package were removed11. Survival analysis was performed for 13 of these datasets, which included 1,581 patients with annotated time to death or last time of follow-up.

### Implementation of Subtype Classifiers

Subtype classifiers were re-implemented in R^12^ using original data as described by Konecny^6^, Verhaak^5^, and Helland^8^. Implementations were validated by reproducing a result from each of the original publications (Supplemental File, Section ‘Reproduction of Published Ovarian Cancer Subtype Classifiers’).

### Survival Analysis

Subtype calls from all included datasets were combined to generate a single Kaplan-Meier plot for each subtyping algorithm (stratified by subtype). Hazard ratios for overall survival between subtypes was estimated by Cox proportional hazards, and statistical significance was assessed by log-rank test, using the *survcomp* R package^13^. Hazard ratios were calculated using the lowest-risk subtype as the baseline group.

### Prediction Strength

Prediction Strength^14^ is defined as a measure of the similarity between pairwise co-memberships of a validation dataset from class labels assigned by (1) a clustering algorithm and (2) a classification algorithm trained on a training dataset (Supplementary Figure 6). The quantity is an established measure of cluster robustness and its interpretation is straightforward: a value of 0 or below indicates poor concordance, and a value of 1 indicates perfect concordance between models specified from training and validation data. Tibshirani and Walther^14^, and subsequent applications of Prediction Strength^15^, have considered a value of at least 0.8 to be an evidence of robust clusters. Prediction Strength was computed as implemented in the *genefu* Bioconductor package^16^.

The tumors in each dataset were clustered *de novo* using our reproduced implementations of the algorithms of Konecny, TCGA/Verhaak, and Tothill (Supplemental File, Section ‘Reproduction of Subtype Clustering Methods’). Each dataset was also classified using implementation of the originally published subtype classifiers. This produced two sets of subtype labels for each sample in each validation dataset; these labels were used to compute Prediction Strength.

### Concordance Analysis

For each pair of classifiers, subtypes were mapped across methods based on concordance, i.e., proportion of patients that were classified as the same (mapped) subtype. Subtype assignment was denoted as concordant only if the pairwise subtype mapping resulted in unique subtypes for each classifier. In other words, if ϕ _ij_ is the mapping of subtypes in classifier *i* to subtypes in classifier *j* (for *i,j* in {1,2,3}), then the following must be satisfied for each subtype : ϕ _12_ (ϕ _23_ (ϕ _31_ (*s*))) = *s*. For the purpose of this study, we considered three methods, each of which strictly classifies the patients into distinct subtypes. For datasets resulting in concordant subtype assignments, patients of the same (mapped) subtype across classifiers were assigned to the subtype names proposed by Verhaak *et al.^5^*. Statistical significance of concordance was assessed by Pearson’s Chi-squared test.

### Filtering tumors by classification margin

Each subtype classifier outputs for each patient a real-valued score for each subtype. Marginally classifiable tumors were identified based on the difference between the top two subtype scores, denoted as the ‘margin’ value. Thus, a higher margin indicates a more confident classification. For each pair of subtype classifiers, classification concordance was assessed on both the full dataset and considering only patients classified with margins above a user-defined cutoff. Concordance was defined as before (proportion of patients that are classified as the same mapped subtype across methods) to calculate concordance between datasets.

### Building a *consensus* classifier

The *consensusOV* classifier was implemented using a Random Forest classifier trained on concordant subtypes across multiple datasets. The Random Forest method has previously been used for building a multi-class consensus classifier to resolve inconsistencies among published colorectal cancer subtyping schemes^17^. In order to avoid merging expression values across datasets, binary gene pair vectors were used as feature space, as recently applied for breast cancer subtyping^18,19^. Since the feature size of this classifier increases quadratically with respect to the size of the original gene set, we used the smallest gene set of the original subtype classifiers (the gene set of Verhaak *et al*.^5^), which contains 100 gene symbols. The *consensusOV* classifier outputs the subtype classification and a real-valued margin score to discriminate between patients that are of well-defined or indeterminate subtype. Similarly to previously published subtype classifiers, a higher margin score indicates higher confidence of classification.

### Leave-one-dataset-out cross-validation

Performance of the consensus classifier for identifying concordantly classified subtypes was assessed using leave-one-dataset-out cross-validation^20^. Concordant subtypes were identified to train the Random Forest classifier using 14 of the 15 datasets, and subtype predictions were tested in the remaining left-out dataset. This process was repeated for all 15 datasets. While predicting the samples in any given dataset, the training set was subsetted to contain only the concordant subtypes in other datasets.

### Correlation analysis with Histopathology and Tumor Purity

Subtype calls from the Consensus Classifier were analysed for correlation with histopathology and tumor purity in the TCGA dataset. In order to best represent the most confident subtype calls, a default cutoff was used to include only the 25% of patients with the largest classification margins. Available histopathology variables included lymphocyte, monocyte, and neutrophil infiltration. Tumor purity was assessed using the ABSOLUTE algorithm^21^, which estimates purity and ploidy from copy number and SNP allele frequency from SNP genotyping arrays (Synapse dataset syn3242754). Significance of associations were tested by one-way ANOVA for patient age, purity, and immune infiltration.

### Research reproducibility

All results are reproducible using R/Bioconductor^22^ and knitr^23^. The code is open source and provided in the *consensusOV* R package (hhttp://www.github.com/bhklab/consensusOV). Literate programming output is provided as a Supplemental File.

## Results

We performed a meta-analysis of three published subtyping algorithms for ovarian cancer ^5,6,8^ and developed a new consensus classifier to identify unambiguously classifiable tumors (Table 1). Each of these algorithms identified four distinct HGS subtypes with specific clinical and tumor pathology characteristics (Figure 1). We assessed the algorithms on a compendium of 15 datasets including over 1,700 HGS patients (Supplemental Table 1) with respect to concordance, robustness, and association to patient outcome. By modifying individual algorithms to discard tumors of intermediate subtype, we found that concordance between algorithms is greatly improved.

**Figure 1:**
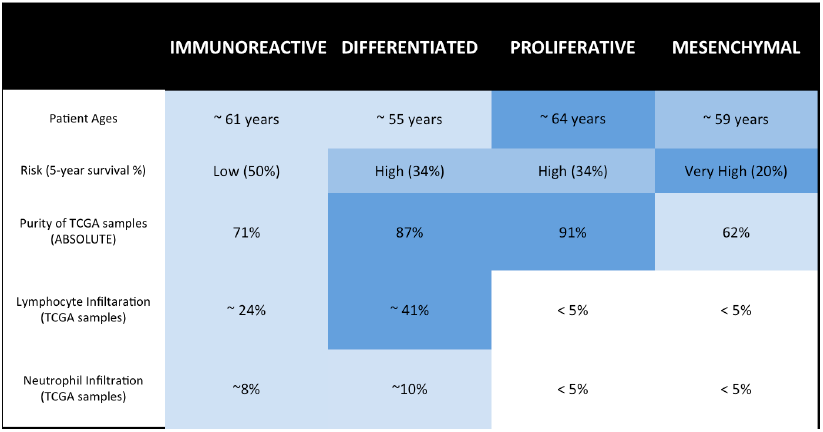
Properties of Subtypes identified by Consensus Classifier. Subtype associations with patient age and overall survival were assessed across our compendium of microarray datasets; association with tumor purity and immune cell infiltration was assessed using the TCGA dataset. Tumor purity was estimated from genotyping data in TCGA; lymphocyte infiltration was based on pathology estimates from TCGA. Patient age (p < 0.001), overall survival (p < 0.005), and ABSOLUTE purity (p < 0.001) were statistically significant across subtypes. When compared to all other groups, the Immunoreactive subtype had elevated infiltration of lymphocytes (p < 0.05) and neutrophils (p < 0.10). Mean monocyte infiltration was less than 5% across all subtypes, and was excluded from this analysis. Classification was performed using default parameters, and mean values of each variable are shown.

### Concordance of published classifiers

We re-implemented three published ovarian subtype classifiers^5,6,8^ (Table 1) and applied these methods to new datasets. We ensured correct implementation of classifiers by reproducing results from the original papers (Supplementary Information). When applied to independent datasets, concordance of the three methods was statistically significant (p < 10^−5^, Chi-square test) with the highest agreement observed for Helland and Konecny subtyping schemes (70.9%), followed by Verhaak and Helland (67.4%) and Verhaak and Konecny (58.9%). Cramer’s V coefficients^24^ indicated a strong association between subtypes as identified by the different algorithms (> 0.5).

### Tumors of intermediate subtype

The individual subtyping algorithms calculate numeric scores for each subtype (mesenchymal, differentiated, immunoreactive, and proliferative), and assign each tumor to the subtype with the highest score. A tumor with a large difference or “margin” between the highest and second highest scores can be considered distinctly classifiable, whereas a tumor with two nearly equal scores could be considered of intermediate subtype. We examined the effect of modifying the individual algorithms to prevent assignment of indeterminate cases at various thresholds. For each pair of subtype classifiers, we examined the classification concordance with increasing thresholds on the margins.

For all pairs of subtype classifier, classification concordance increased as additional marginal cases are removed, approaching 100% concordance once the majority of tumors are left unclassified (Figure 2B). Three-way concordance followed the same trend with lower overall concordance: a minimum of 23% for the proliferative subtype and maximum of 45% for the immunoreactive subtype when all tumors are classified. Restricting the concordance analysis to the top 50% of tumors by margin value resulted in an increased overlap between 35% (proliferative) and 65% (immunoreactive). At a strict threshold of where only 10% of tumors are classified, 88% of tumors overall are concordantly classified by all three published subtyping algorithms (Figure 2C). This large gain in concordance results from large reductions in both singleton calls - tumors assigned to one subtype by one algorithm, but not by the other two algorithms - and in 2-to-1 calls, tumors assigned to one subtype by two algorithms, but not by the third (Figure 2D). This indicates that tumors distinctly classifiable by a single algorithm are more likely to be concordantly classified by the other algorithms, and conversely, tumors that appear ambiguous to one algorithm are less likely to be classified in the same way by the other algorithms.

**Figure 2:**
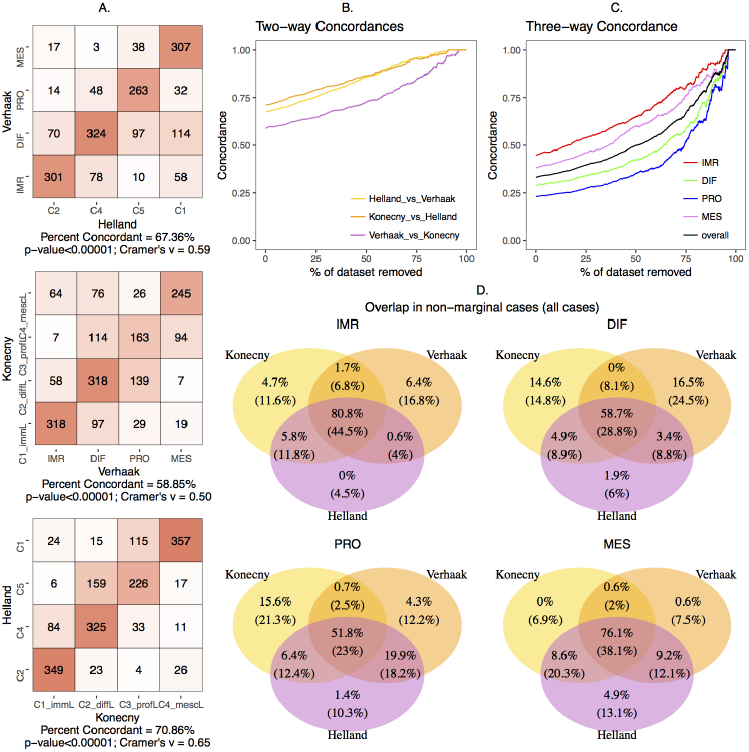
Concordance Analysis. (A) Contingency table showing concordance of subtypes while comparing the methods pairwise (B) Pairwise concordance between the methods versus percentage of the dataset with samples of lower subtype margins removed, (C) three-way overall concordance between the methods and that of the individual subtypes versus percentage removed, (D) The classification of patients by three published algorithms as a Venn diagram for each of the four subtypes. Each area shows percentages of patients when all patients are classified (below, in parentheses) and after refusing to classify 75% of the most marginally classified tumors by any of the three methods (above). Thus, the numbers on the top of the three-way intersection are the concordant tumors according to the three original algorithms. Bottom numbers indicate relatively unambiguous subtype predictions by all three algorithms and which are also concordant with the others.

### Survival Analysis

All proposed subtyping algorithms classified patients into groups that significantly differed in overall survival (Figure 3A, p < 10^−5^ for each subtyping algorithm, log-rank test). Comparing low-risk to high-risk subtypes for each algorithm, the hazard ratios increase from approximately 1.5 as marginal cases are removed (Figure 3B), suggesting that marginal cases may contribute to the intermediate survival profiles between subtypes.

**Figure 3:**
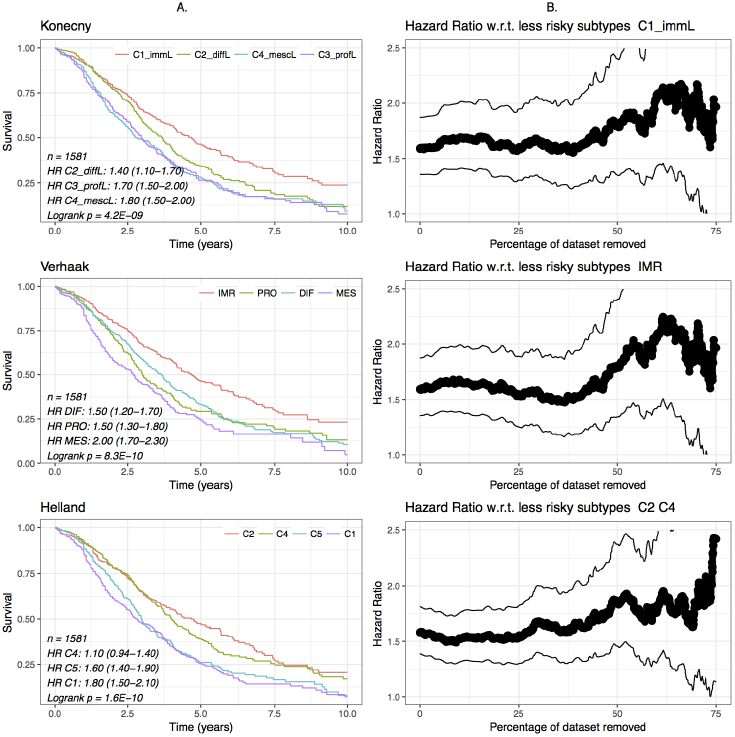
Survival Analysis. (A) Kaplan-Meier curves of subtypes of the 1581 patients with survival data under different methods. (B) Hazard ratios and 95% confidence intervals of the lowest-risk subtype (Konecny and Verhaak) or two subtypes (Helland) compared to the remaining subtypes.

### Robustness of the Classifiers

Robust molecular subtyping should be replicable in multiple datasets. We performed *de novo* clustering in 15 independent ovarian datasets using the authors’ original gene lists and clustering methods. We compared these *de novo* clusters to the labels from our implementation of the published classifiers to assess robustness using the Prediction Strength (PS) statistic^14^. For PS estimation, we included validation datasets with at least 100 HGS tumors. Overall we observed low robustness for all classifiers, with PS values under 0.6 for the three algorithms across datasets (Supplementary Figure 7), none meeting the 0.8 threshold typically indicating robust classes^14,15^.

To assess whether low confidence predictions are driving the PS estimation, we re-computed the robustness of each algorithm set to classify varying fractions of the tumors with the highest margins. We used the largest dataset available, the TCGA dataset, as the validation set, and varied margin cutoffs of the Tothill and Konecny classifiers to require them to classify between 25% and 100% of the cases. From 10 random clustering runs, we report the median PS for the dataset. Clustering was performed on the full TCGA dataset and tumors of low margin values were removed subsequent to clustering and after the classifier was fully defined, in order to avoid optimistically biasing the apparent strength of clusters. We observed that the robustness of each algorithm is substantially improved by allowing them to refuse to classify ambiguous cases. The Tothill algorithm achieved almost perfect robustness (PS = 0.96) when allowed to leave 75% of cases unclassified (Figure 4).

**Figure 4:**
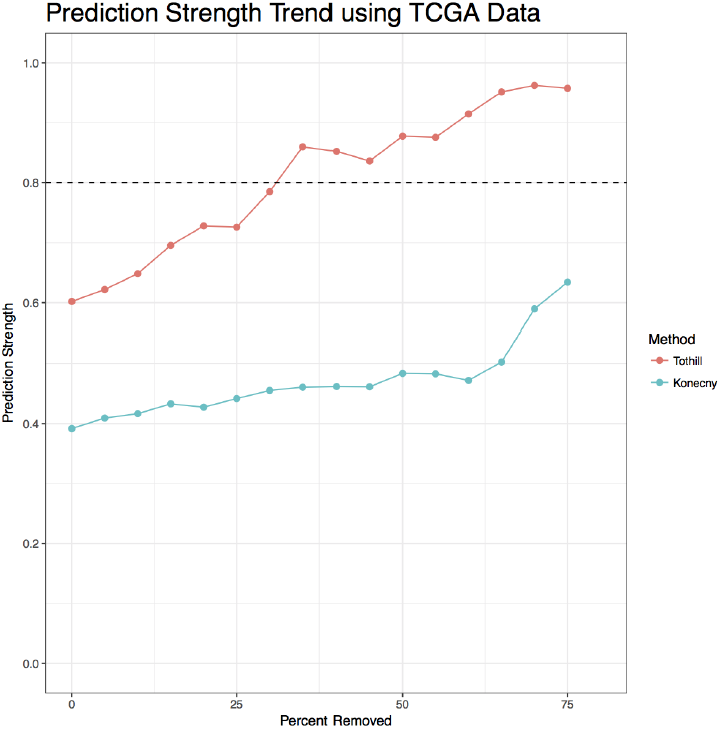
Robustness Analysis of published classifiers, by Prediction Strength. In each dataset, concordance was calculated between the published classifier and a classifier re-trained on the validation dataset. The TCGA dataset also classified using the published classifiers of Helland and Konecny (no re-training was done for the classifiers). The TCGA dataset was also clustered using the methods of Tothill and Konency (in red and blue respectively). Samples were removed from Prediction Strength calculations starting with the most ambiguous samples (with the smallest difference between the top subtype prediction and runner-up subtype prediction); the x-axis shows the percent removed before computing prediction strength. Each algorithm improves in robustness when allowed to leave ambiguous samples, that it is less certain in its classification, unclassified.

### Consensus Classifier

To maximize concordance across classifiers, we developed *consensusOV*, a consensus subtyping scheme facilitating classification of tumors of well-defined subtypes (Figure 5). This classifier uses binary gene pairs^18,19^ to support application across gene expression platforms. The *consensusOV* classifier exhibits overall pairwise concordance of 67 - 78% with each of the other three algorithms, when classifying all tumors; and 94% concordance with tumors that are concordantly classified by the other three algorithms (Figure 5A). The margins of *consensusOV* are higher for concordantly classified cases than for non-concordantly classified cases, and this difference in margins is greater than for any of the other three classifiers (Figure 6A). Accordingly, *consensusOV* was also most effective in identifying concordantly classified cases, although it was similar to the Konecny classifier in this respect (AUC = 0.76, Figure 6B). As expected, differences in survival of subsets identified by *consensusOV* are similar to those identified by previous classifiers. The highest risk subtypes are proliferative (HR=1.44, 95% CI: 1.07-1.94) and mesenchymal (HR=1.97, 95% CI: 1.46-2.67) when removing 75% of indeterminate low-margin tumors, with similar hazard ratios for the concordant cases (Figure 5B).

**Figure 5:**
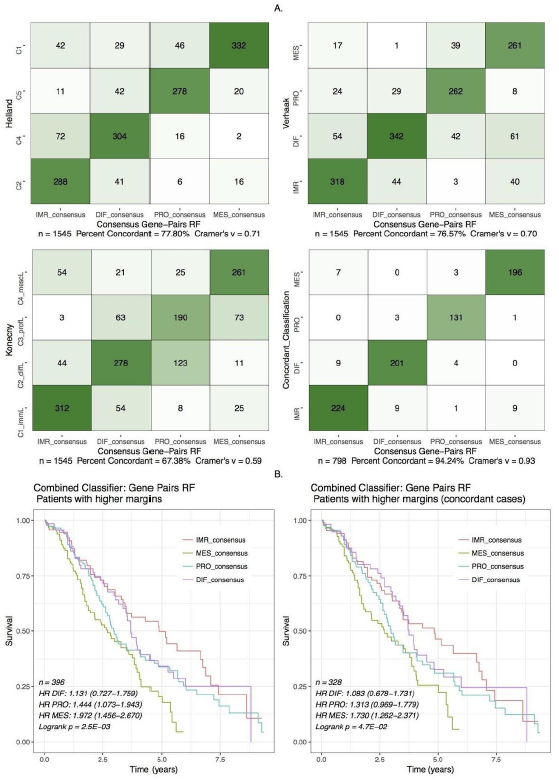
Concordance and Survival Stratification of consensusOV. (A) Contingency plots showing concordance of subtype classification between *consensusOV* and the classifiers of Helland, Verhaak, Konecny. To address differences in gene expression scales due to different experimental protocols, *consensusOV* standardizes genes in each dataset to the same mean and variance, and computes binary gene pairs. The fourth (bottom-right) plot shows the concordance between the consensus classifier and the patients concordantly classified between the three classifiers. (B) Survival curves for the pooled dataset provided by *consensusOV*. Classification was performed using leave-one-dataset-out validation. For the bottom two figures, classification with *consensusOV* was performed with the default cutoff, in which 75% of patients with the lowest margin are not classified.

**Figure 6:**
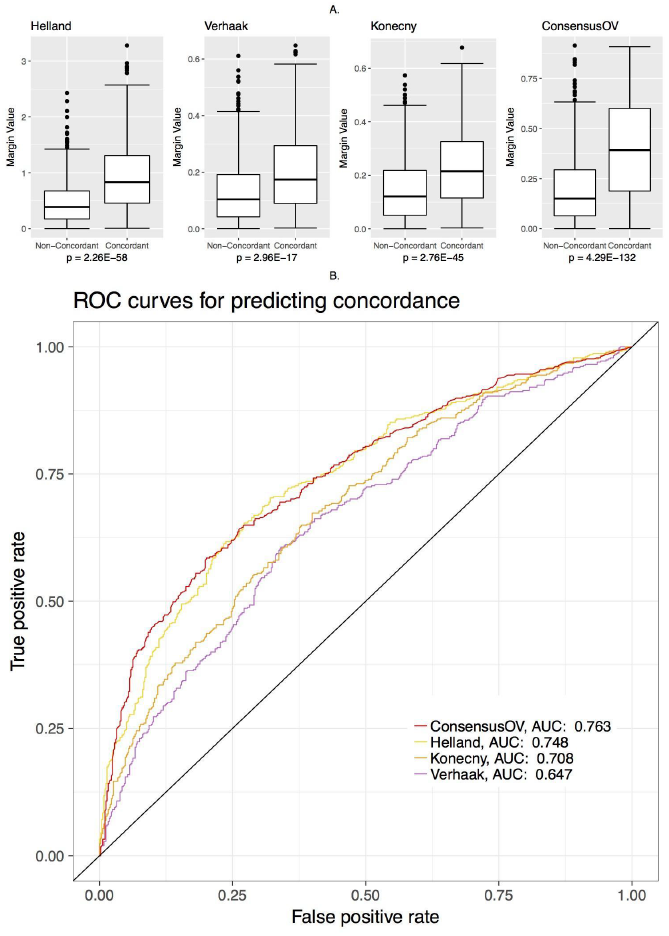
Margin Analysis. (A) Boxplots indicating the margin values assigned by each classifier to concordant and discordant cases. All statistical tests were performed using the Wilcoxon rank-sum test. (B) ROC curve for assessing the ability of margin values to discriminate between concordant and discordant cases.

## Discussion

The existence of four distinct and concordant molecular subtypes of HGOC has been reported in several studies of large patient cohorts^4–6,8^, but also called into question by another effort^2^ that could not identify subtypes, and by an independent validation effort that reported only two or three reproducible subtypes^25^. Meanwhile, significant effort is being expended to translate these subtypes to clinical practice, for example to predict response to the angiogenesis inhibitor bevacizumab in the ICON7 trial^26,27^. This article pursues three major objectives: (1) reproduction of published subtype classification algorithms as an easily usable open-source resource; (2) evaluation of the robustness and prognostic value of each proposed subtyping scheme in independent data; and (3) consolidation of proposed subtyping schemes into a consensus algorithm.

We find that while the proposed 4-subtype classifications demonstrate significant concordance and association with patient survival, none are robust to re-training in new datasets. By modifying any of these algorithms to refuse to identify tumors of ambiguous subtype, robustness and concordance of subtyping algorithms improve dramatically. We propose a “consensus” classifier for 25% of tumors that can be classified with high confidence regardless of training dataset, although a continuous trade-off exists between classifying more tumors versus having greater confidence in those classified.

Ambiguity in tumor classification might arise from a heterogeneous admixture of different subtypes, or from a more homogeneous composition of indeterminate subtype. This distinction has implications for the therapeutic value of the proposed subtypes. Lohr et al. estimated that 90% of tumors in the TCGA HGS dataset are polyclonal^28^, and clonal spread of HGS ovarian cancer has been directly inferred from single-nucleus sequencing^29^. However, it remains unclear whether multiple clones in a tumor are consistently classifiable to the same subtype. If a tumor consists of multiple clones of different subtypes, then a subtype-specific therapy will likely lead to relapse as other clones survive and continue to grow. If this situation is common, even unambiguously classifiable tumors might be contaminated by small amounts of another subtype that could lead to relapse after subtype-specific therapy. This question could not be resolved by the current datasets, but may eventually be addressed by single-cell RNA sequencing^30^ which is expected to further improve precision HGS molecular subtyping.

Several findings stand out in the validation of published subtyping algorithms. First, although previous studies reported inconsistent findings on whether subtypes differ by patient survival, our analysis in independent data showed clear survival differences. The 5-year survival rate for patients with different subtypes ranged from as low as 20% to as high as 50%. Second, published algorithms do not meet previously defined standards of robustness in terms of Prediction Strength, a measure of consistency between subtype classifiers trained in independent datasets. Finally, the concordance of three algorithms, established independently by different research groups from different patient cohorts, is only moderate but can be greatly improved by modifying the original algorithms to allow them to leave ambiguous tumors unclassified. In their original forms, all-way concordance of the four defined classes occurs in 23% to 45% of tumors. As the individual algorithms are modified to leave ambiguous cases unclassified, the minority of tumors remaining can be classified with over 90% concordance between the three algorithms. Unfortunately, these ambiguous cases account for up to 75% of HGSC of the ovary. This has important implications for the clinical application of subtypes in ovarian cancer.

Moving forward, general agreement on how molecular subgroups are defined is expected to facilitate the use of expression data in clinical trial design, thereby improving prognosis as well as treatment benefit^31^. We introduced the subtype classifier *consensusOV,* which represents the consensus of published HGS subtype classifiers. By training on multiple datasets, using binary (pairwise greater-than or less-than) relationships between pairs of genes, and using a relatively small gene set, *consensusOV* is designed to be applicable across gene expression platforms and datasets, making it effective for benchmarking and meta-analysis. The algorithm provides subtype-specific scores, margin values, and an option to leave tumors of intermediate subtype unclassified. Identifying tumors of distinct subtype is an important step towards defining robust and therapeutic phenotypes.

## Funding

G.M. Chen was supported by the Canadian Institutes of Health Research and the Terry Fox Research Institutes. D.A.M. Gendoo was supported by the Ontario Institute for Cancer Research through funding provided by the Government of Ontario. Z. Safikhani was supported by The Cancer Research Society (Canada). B. Haibe-Kains was supported by the Gattuso-Slaight Personalized Cancer Medicine Fund at Princess Margaret Cancer Centre, the Canadian Institutes of Health Research, and the Terry Fox Research Institutes. L Waldron was supported by grants from the National Cancer Institute at the National Institutes of Health (1R03CA191447-01A1 and U24CA180996).

### Acknowledgements

The authors thank Brad Nelson for his feedback regarding the prognostic value of molecular subtypes in ovarian cancer, and Andrew Cherniak for providing ABSOLUTE purity and ploidy estimates for tumors from The Cancer Genome Atlas.

